# Absence of receptor guanylyl cyclase C enhances ileal damage and reduces cytokine and antimicrobial peptide production during oral *Salmonella* Typhimurium infection

**DOI:** 10.1101/214163

**Authors:** Shamik Majumdar, Vishwas Mishra, Somesh Nandi, Mudabir Abdullah, Anaxee Barman, Abinaya Raghavan, Dipankar Nandi, Sandhya S. Visweswariah

**Author notes:** Address correspondence to Sandhya S. Visweswariah. SM, VM and SN contributed equally to this work.

## Abstract

Non-typhoidal *Salmonella* disease contributes towards significant morbidity and mortality across the world. Host factors including IFN-γ, TNF-α and gut microbiota, significantly influence the outcome of *Salmonella* pathogenesis. However, the entire repertoire of host protective mechanisms contributing to *Salmonella* pathogenicity is not completely appreciated. Here, we have investigated the roles of receptor guanylyl cyclase C (GC-C) that is predominantly expressed in the intestine, and regulates intestinal cell proliferation and fluid-ion homeostasis. Mice deficient in GC-C (*Gucy2c*^-/-^) displayed accelerated mortality following infection via the oral route, in spite of possessing comparative systemic *Salmonella* infection burden. Survival following intra-peritoneal infection remained similar, indicating that GC-C offered protection via a gut-mediated response. Serum cortisol was higher in *Gucy2c*^-/-^ mice, in comparison to wild type (*Gucy2c*^+/+^) mice, and an increase in infection-induced thymic atrophy, with loss in immature CD4^+^CD8^+^ double positive thymocytes, was observed. Accelerated and enhanced damage in the ileum, including submucosal edema, epithelial cell damage, focal tufting and distortion of villus architecture, was seen in *Gucy2c*^-/-^ mice, concomitant with a larger number of ileal tissue-associated bacteria. Transcription of key mediators in *Salmonella*-induced inflammation (IL-22/Reg3β) were altered in *Gucy2c*^-/-^ mice in comparison to *Gucy2c*^+/+^ mice. A reduction in fecal Lactobacilli, which are protective against Salmonella infection, was observed in *Gucy2c*^-/-^ mice. *Gucy2c*^-/-^ mice cohoused with wild type mice continued to show reduced Lactobacilli and increased susceptibility to infection. Our study therefore suggests that receptor GC-C confers a survival advantage during gut-mediated *S*. Typhimurium pathogenesis, presumably by regulating *Salmonella-effector* mechanisms and maintaining a beneficial microbiome.

## Introduction

Salmonellosis, the disease caused by the Gram-negative intracellular bacterium, *Salmonella* manifests in humans as typhoid, paratyphoid fever, non-typhoidal septicemia or gastroenteritis. Infection with *Salmonella* Typhi and Paratyphi serotypes are limited to humans, and cause the systemic disease, typhoid fever. Infections with other serotypes, called non-typhoidal *Salmonella,* result in enteritis and diarrhoea. *Salmonella* infects hosts orally via contaminated food or water (1). Mice infection models are useful not only to study factors in the pathogen that regulate disease progression, but also host factors that can modulate the severity of response to *Salmonella* infection (2). In susceptible mouse strains, such as those with defects in the gene encoding Slc11a1 (Nramp1), the bacterium traverses the distal ileum, and mice develop systemic infection after colonizing the Peyer’s patches, liver and spleen (3).

Colonization of the mouse intestine is dependent on a number of factors, that include commensal microbiota, the gut mucosa and the gut-associated immune system. A number of host factors such as IFN-*γ* (4), TNF-*α* (5, 6) and IL-22 modulate *Salmonella* pathogenesis (1). These factors regulate the production of chemokines that recruit inflammatory cells to the site of tissue damage and infection (7). In addition, it is conceivable that modulators of intestinal epithelial cell function may also provide mechanisms to modulate *Salmonella* pathogenesis, since the first point of contact for the pathogen is the epithelial cells that line the gastrointestinal tract (8).

Receptor guanylyl cyclase C (GC-C) is a transmembrane receptor predominantly expressed on the apical membrane of the intestinal enterocytes. It serves as a receptor for the endogenous hormones, guanylin and uroguanylin (9, 10). Activation of GC-C upon ligand-binding results in production of cGMP, which leads to protein kinase G II-dependent activation of the cystic fibrosis transmembrane conductance regulator (CFTR) and inhibition of NHE3, the sodium–hydrogen exchanger 3. Activation of CFTR causes efflux of chloride and CFTR-dependent bicarbonate secretion into the lumen and inhibition of sodium absorption by NHE3, thus resulting in an osmotic gradient, leading to fluid accumulation in the lumen. Hence, the major role of GC-C is in maintenance of fluid-ion homeostasis in the gut (11). GC-C also serves as the receptor for the heat-stable enterotoxin (ST), the causative agent of enterotoxigenic *Escherichia coli* (ETEC)-mediated diarrhoea in children and traveller’s diarrhoea (12). Mice lacking GC-C are resistant to heat-stable enterotoxin-induced fluid accumulation (13), and are reported to display higher susceptibility to dextran sodium sulphate-induced colitis (14). Previously, we have demonstrated that mice lacking GC-C show increased N-Methyl-N-nitrosourea-induced aberrant crypt foci (15), indicating a role for GC-C in maintaining intestinal cell proliferation.

Mice deficient in GC-C (i.e., *Gucy2c*^-/-^) mice have been reported to possess a compromised gut epithelial barrier which was attributed to the lower expression of tight junction proteins such as occludin, claudin 2 and claudin 4 (14). *Citrobacter rodentium* (*C*. *rodentium*), an enteric Gram-negative bacterium causes self-limiting infection in mice, and *Gucy2c*^-/-^ mice are reported to be more susceptible to infection by them (16). Here, we have studied the role of GC-C as a host factor during *Salmonella*-mediated pathogenesis in mice. We find that in contrast to expectations that a compromised gut barrier would lead to enhanced systemic infection, the increased susceptibility of *Gucy2c*^-/-^ mice to oral infection with *Salmonella* Typhimurium (S. Typhimurium; St) was a result of increased bacterial load in the ileum, altered cytokine production, increased ileal tissue damage, and coupled to an altered fecal microbiome.

## Results

### Mice lacking GC-C display accelerated mortality during oral *S*. Typhimurium infection

We infected C57BL/6 wild type (*Gucy2c*^+/+^) and *Gucy2c*^-/-^ mice orally with *S*. Typhimurium and monitored their survival. We chose not to treat the mice with streptomycin (17), since we were interested in an acute model of *S*. Typhimurium infection, without altering the existing microbiota in these mice (18). Surprisingly, survival experiments revealed an increased susceptibility of *Gucy2c*^-/-^ mice to oral, but not intra-peritoneal, infection with *S*. Typhimurium (Fig. 1a), indicating a role for GC-C in offering protection to gut-associated infection. Median survival for *Gucy2c*^+/+^ mice following oral infection was 112 h while it was 80 h for *Gucy2c*^-/-^ mice, and the difference was statistically significant (*p* = 0.006). Interestingly, fecal excretion of bacteria was similar in *Gucy2c*^+/+^ and *Gucy2c*^-/-^ mice at early stages of infection, as was the bacterial burden in organs such as the spleen, Peyer’s patches and liver on day 3 post infection (Fig. 1b, c). In agreement with earlier observations (19), poor survival on *Salmonella* infection was correlated with a significant reduction in body weight of *Gucy2c*^-/-^ mice on day 3 (Fig. 1d).

**Figure 1:**
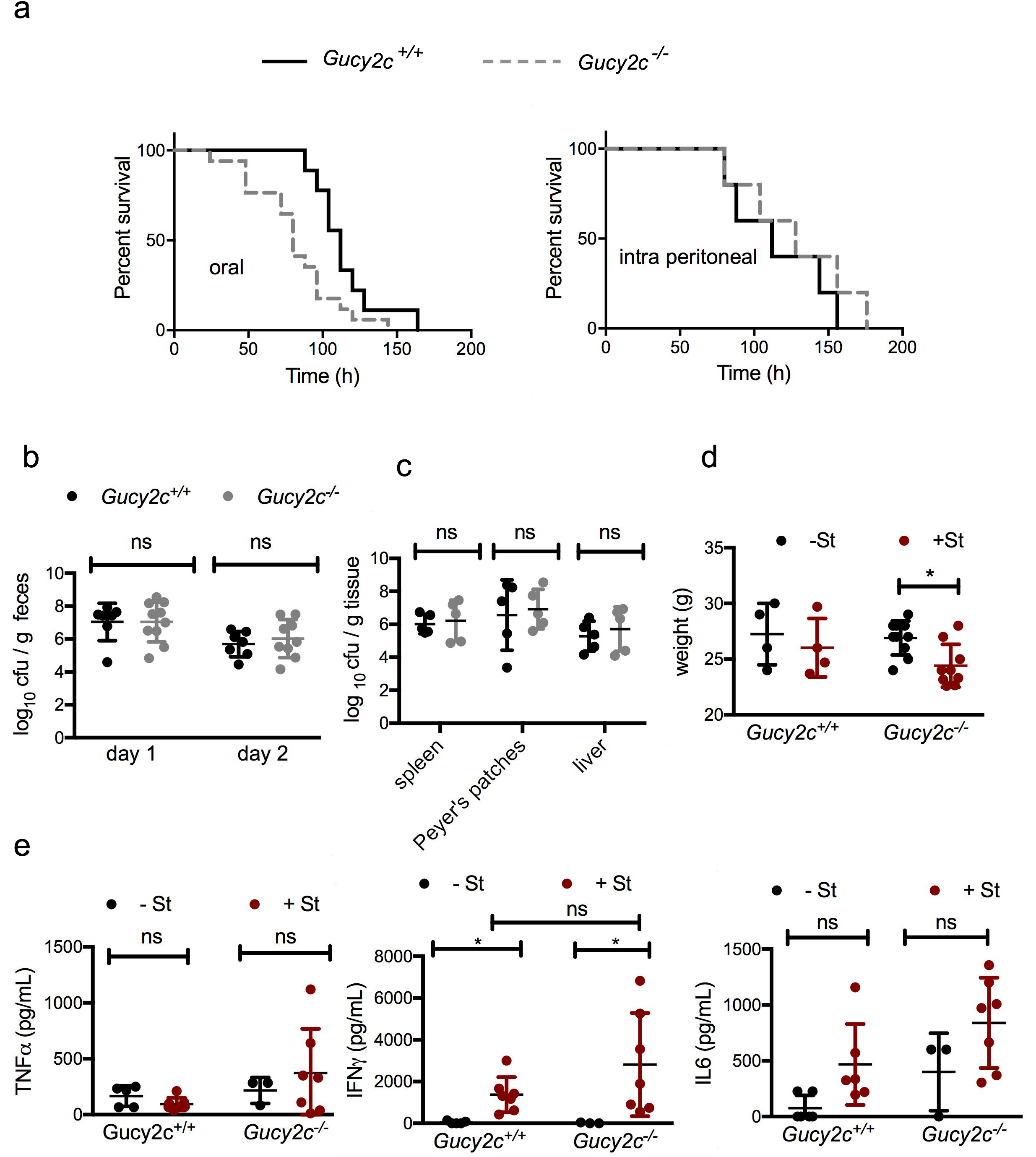
*Gucy2c*^-/-^ mice display accelerated mortality during oral infection with *S*. Typhimurium. (a) *Gucy2c*^+/+^ and *Gucy2c*^-/-^ mice were infected with *S*. Typhimuirum either orally or intra-peritoneally. The mice were monitored for survival at 8 h intervals. Survival curves for oral and intra-peritoneal infection were obtained from independent experiments containing a total of 10-11 mice. (b) Mice (*Gucy2c*^+/+^, black circles; *Gucy2c*^-/-^, grey circles) were orally infected with *S*. Typhimurium and the bacterial load in the feces on day 1 and 2 following infection were estimated. Data show values for individual mice, with error bars showing the mean ± SD. (c) Infected mice were sacrificed on day 3, tissues harvested and bacterial burden in the spleen, Peyer’s patches and liver was estimated. Data show values for individual mice, with error bars showing the mean ± SD. (d) Mice were weighed prior to oral infection and 3 days post infection. (e) Serum was collected from uninfected and infected mice and levels of TNF-α, IFN-γ and IL-6 were measured by ELISA. In panels b-e, the data are depicted as mean ± SD of 3-7 mice per group. Statistical significance among the experimental groups in all panels was analysed using two-way ANOVA, \**p*≤0.05 and ns: not significant.

Next, we monitored pro-inflammatory cytokines in the sera 3 days post oral infection, to determine whether enhanced immune cell activation and inflammation was responsible for the poor survival of *Gucy2c*^-/-^ mice. As shown in Fig. 1e, while there was a tendency for an increase in serum TNF-*α* and IL-6 levels on infection, the increase was not statistically significant. IFN-*γ* levels were elevated on infection in both strains of mice, but there was no difference between Gucy2c^+/+^ and *Gucy2c*^-/-^ mice. Therefore, we conclude that an increase in systemic infection or inflammation was not the cause for early death of *Gucy2c*^-/-^ mice.

### Enhanced infection-induced thymic atrophy is observed in *Gucy2c*^-/-^ mice

We have previously reported that serum cortisol, a marker of general stress experienced by the animal, is increased on *Salmonella* infection and contributes towards *S*. Typhimurium-induced thymic atrophy in mice (20, 21). We monitored cortisol levels and observed an increase in serum cortisol on infection. However, at 48 h, *Gucy2c*^-/-^ mice showed higher levels of circulating cortisol than that seen in *Gucy2c*^+/+^ mice (Fig. 2a). Thymic atrophy was observed in both *Gucy2c*^+/+^ and *Gucy2c*^-/-^ mice (Fig. 2b, c), but was more pronounced in *Gucy2c*^-/-^ mice, both at 48 and 96 h post infection (Fig. 2b). CD4 and CD8 cell surface staining of isolated thymocytes revealed an increased depletion of CD4^+^CD8^+^ thymocytes in *Gucy2c*^-/-^ mice post 96 hrs of infection (Fig. 2c).

**Figure 2.**
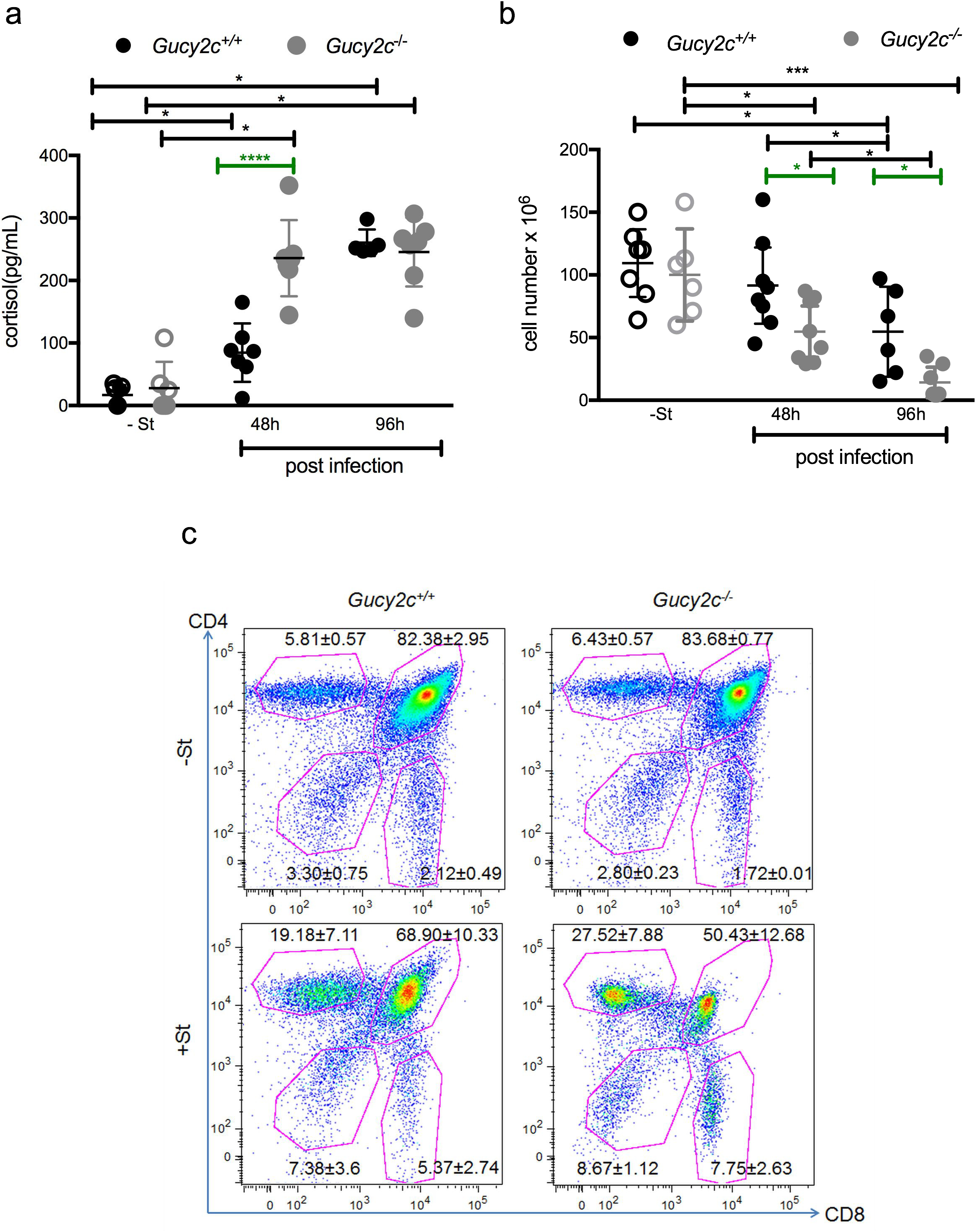
Early rise in serum cortisol amounts and increased infection-induced thymic atrophy is observed in *Gucy2c*^-/-^ mice. (a) *Gucy2c*^+/+^ and *Gucy2c*^-/-^ mice were orally infected with *S*. Typhimurium. At the indicated time points, uninfected (-St) and infected (+St) mice were sacrificed and serum cortisol levels were measured by ELISA. Data show values for individual mice, with error bars showing the mean ± SD. (b) Viable cells from harvested thymi were quantified by Trypan blue exclusion assay using a haemocytometer. (c) The thymocytes from uninfected (-St) and mice post 96 h of infection (+St) were stained for cell surface expression of CD4 and CD8 and the density plots were constructed to quantify the percentages of the major cell populations. Data are depicted as mean ± SEM of 5-8 mice per group. In all panels, experimental groups were analysed for statistical significance using two-way ANOVA, \**p* ≤ 0.05, \*\*\**p* ≤ 0.001 and \*\*\*\**p* ≤ 0.0001. Green bars indicate differences that were statistically significant between *Gucy2c*^+/+^ (black circles) and *Gucy2c*^-/-^ (grey circles) mice.

### Increased epithelial damage in the small intestine is observed in *Gucy2c*^-/-^ mice

Results so far suggest that the protective effects of GC-C are mediated predominantly at the level of the gastrointestinal tract, and could be initiated early during infection, thereby resulting in significant weight loss and earlier death in *Gucy2c*^+/+^ mice. We therefore monitored bacterial load in various regions of the gut following infection in both *Gucy2c*^+/+^ and *Gucy2c*^-/-^ mice. Colony forming units (CFU) were enumerated on day 3 following infection, at which time a few *Gucy2c*^-/-^ mice had already succumbed. As shown in Fig. 3a, while bacterial loads in the cecum and colon were similar in both strains of mice, and an almost 2-log order increase in bacteria was seen in the distal ileum.

**Figure 3:**
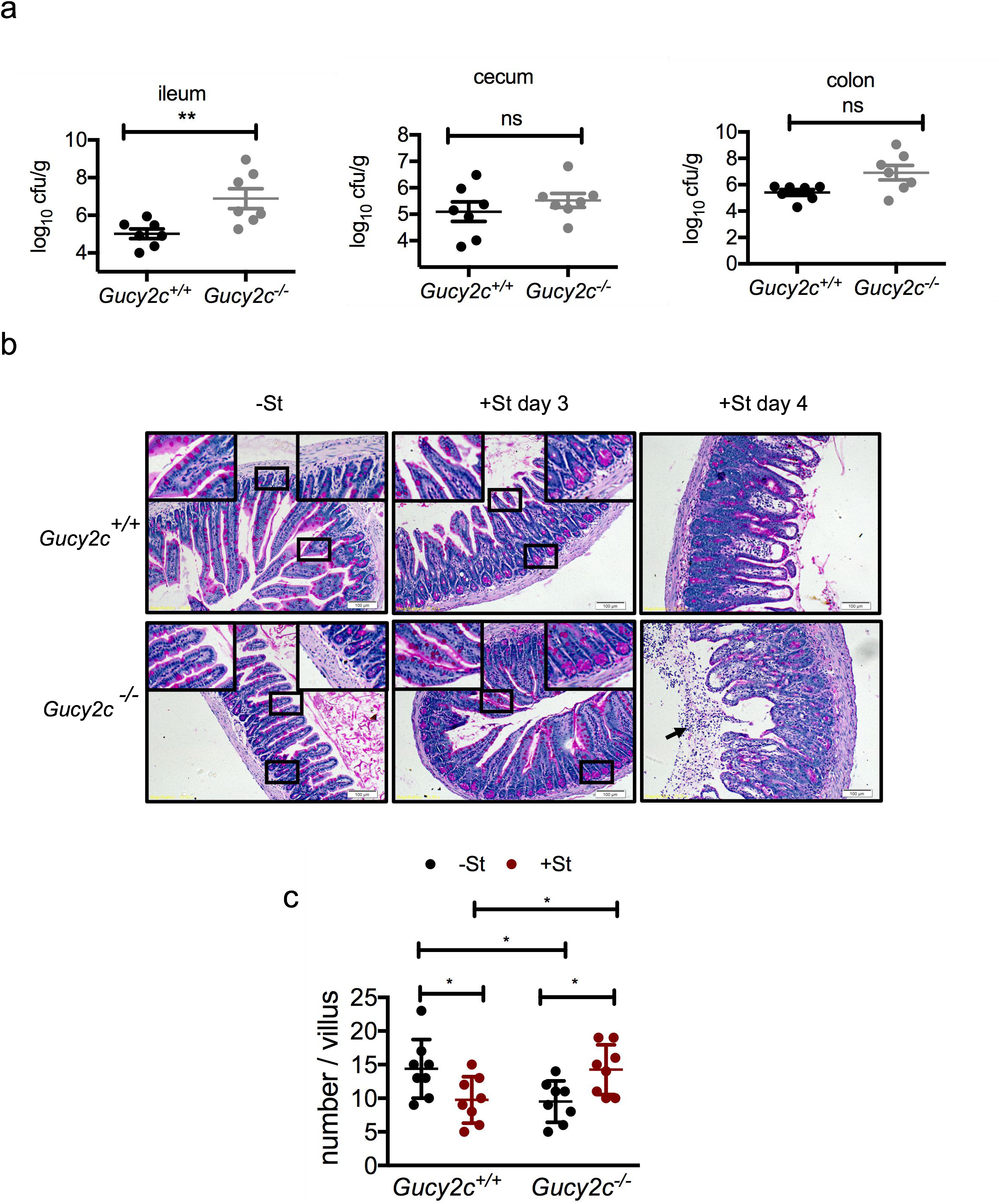
*Gucy2c*^-/-^ mice show increased bacterial colonization and intestinal damage. Infected *Gucy2c*^+/+^ (black circles) and *Gucy2c*^-/-^ (grey circles) mice were sacrificed 3 days post oral *S.* Typhimurium infection. (a) Bacterial load in the ileum, cecum and colon was estimated. Values represent individual mice, and lines show the mean ± SD. (b) Periodic Acid Schiff (PAS) staining was performed on formalin fixed ileal tissue sections obtained from uninfected (-St) and infected (+St) on day 3 and 4 following infection. One representative ileal tissue section from each experimental group containing 3–4 mice is depicted, and the scale bars measure 100 μm. Arrow shows a large number of apoptic cells in the lumen of *Gucy2c*^-/-^ mice. (c) The number of goblet cells was quantified in individual, clearly identifiable, villi in each section from uninfected (−St; black circles) and infected (+St; red circles) mice. Data shown is from individual villi across a section, and lines indicate the mean ± SD. Statistical analysis was performed by two-way ANOVA and \**p* ≤ 0.05.

We measured ileal damage and goblet cell number on day 3, using Periodic Acid Schiff (PAS) staining following paraformaldehyde fixation. Examination of small intestinal sections showed marked histopathological differences between infected *Gucy2c*^+/+^ and *Gucy2c*^-/-^ mice. Infection resulted in more severe inflammation in the small intestine of *Gucy2c*^-/-^ mice, with increased focal tufting and distortion of the crypt-villus architecture by day 4 (Fig. 3b). A larger number of dead cells were also observed in the lumen of the ileum in *Gucy2c*^-/-^ mice by day 4 post-infection.

PAS staining revealed a reduced number of goblet cells in uninfected *Gucy2c*^-/-^ mice (Fig. 3c). Goblet cell numbers reduced in *Gucy2c*^+/+^ on infection, as has been reported earlier (17), but were increased in *Gucy2c*^-/-^ mice. Also noteworthy is the intense staining of Paneth cells in the crypts of both *Gucy2c*^+/+^ and *Gucy2c*^-/-^ mice following infection.

### *S.* Typhimurium infection downregulates the transcription of GC-C and its ligands, guanylin and uroguanylin

Results so far indicate that signaling via GC-C protects the host during *Salmonella* infection, by reducing the extent of tissue damage and colonization in the ileum. We therefore asked if infection alters the expression of genes in the GC-C signaling pathway, which could be a strategy used by *Salmonella* to increase its virulence in the host. We performed quantitative real-time PCR analysis of cDNA prepared from the distal ileum of *Gucy2c*^+/+^ mice on day 3 following oral infection and observed that the transcript levels of GC-C were significantly reduced on infection (Fig. 4). The distal ileum may be a region where both guanylin and uroguanylin are expressed to comparable levels (22), and interestingly, transcript levels of both the ligands were reduced on infection in both *Gucy2c*^+/+^ and *Gucy2c*^-/-^ mice (Fig. 4). Other genes downstream of GC-C did not show a change in expression levels (*Cftr*, *PkgII* and *Pde5*) on infection, while *Nhe3* transcripts were upregulated in both *Gucy2c*^+/+^ and *Gucy2c*^-/-^ infected mice. Activation of GC-C results in cGMP-dependent phosphorylation of NHE3 that inhibits this Na^+^/H^+^ exchanger (11, 23). Therefore, the downregulation of guanylin and uroguanylin transcripts, coupled with increased expression of *Nhe3* would enhance Na+ uptake by the epithelial cells, possibly counteracting loss of this ion during *Salmonella*-induced diarrhea in humans.

**Figure 4.**
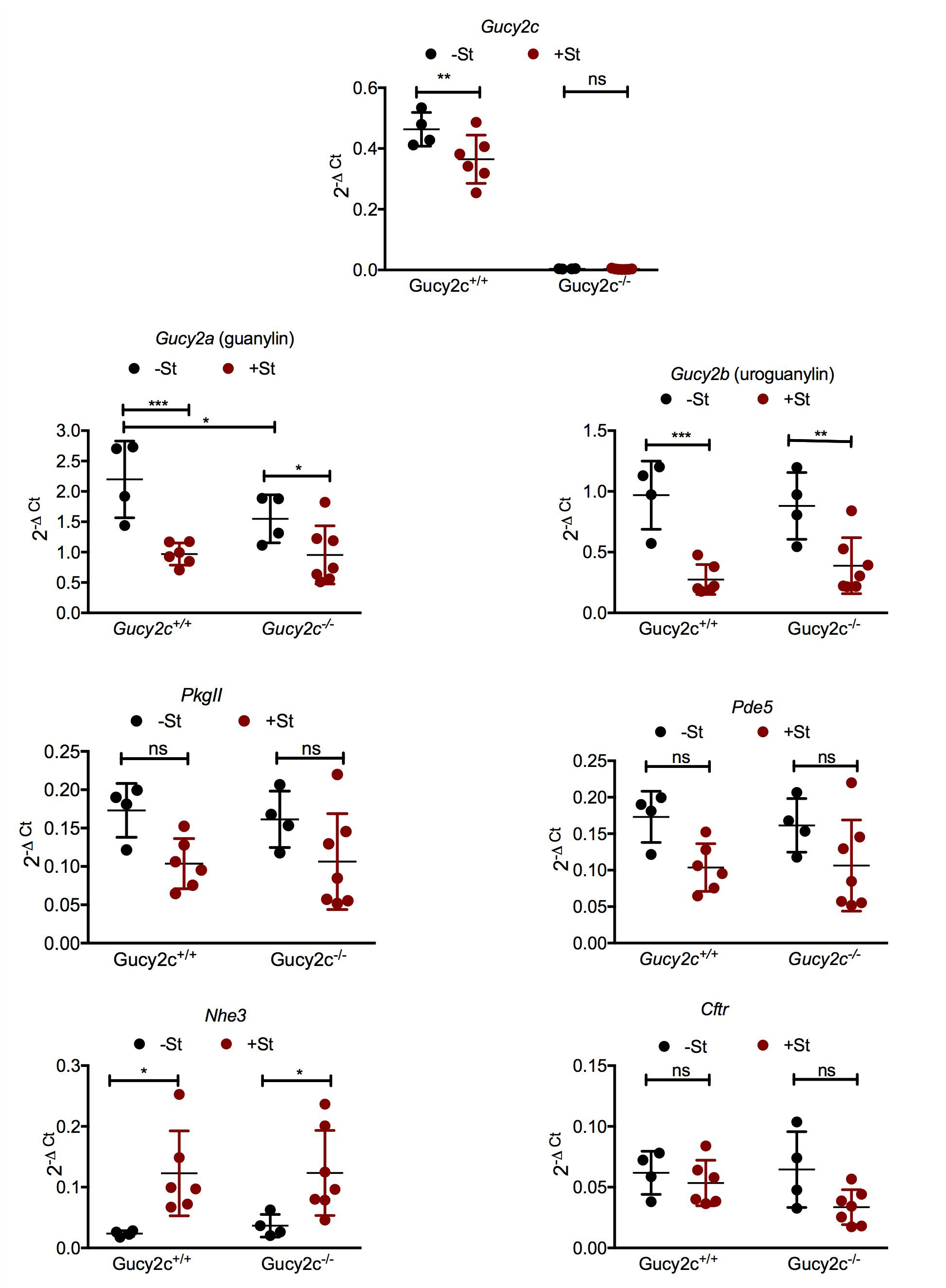
*S.* Typhimurium infection regulates expression of genes in the GC-C signaling pathway. RNA was prepared from ileal tissue of uninfected (-St) and infected (+St) mice on day 3 following oral infection. Quantitative real-time PCR analysis was performed to estimate the transcript levels of *Gucy2c*, *Guca2a, Guca2b, PkgII, Pde5, Nhe3 and Cftr.* The data are depicted as values for individual mice, and lines represent the mean ± SD. Two-way ANOVA was used to analyse for statistical significance among the experimental groups, \**p* ≤ 0.05, \*\**p* ≤ 0.01, \*\*\**p* ≤ 0.001 and ns: not significant.

### GC-C regulates transcription of cytokines and effectors during *Salmonella* infection

Intestinal inflammation following *Salmonella* infection is a consequence of upregulation of pro-inflammatory genes including *TNF-α*, *Il-1β*, *Il-22*, *Il-6*, *Ifn-γ* (24) and genes expressing the C-type lectins *Reg3γ* and *Reg3β* (1). Transcripts of most cytokines were upregulated to similar extents in both *Gucy2c*^+/+^ and *Gucy2c*^-/-^ mice following infection (Fig. 5). Transcript levels of only a few genes (*Il-22*, *Cxcl15* and *Reg3β* ; see green bars in Fig. 5) differed statistically between *Gucy2c*^+/+^ and *Gucy2c*^-/-^ mice following infection. While *lipocalin 2* transcripts were significantly increased in *Gucy2c*^-/-^ mice, there was no statistical difference between levels in wild type and *Gucy2c*^-/-^ mice following infection. In summary, this indicates a role for GC-C signaling in modulating specific arms of the *Salmonella-induced* immune response.

**Figure 5.**
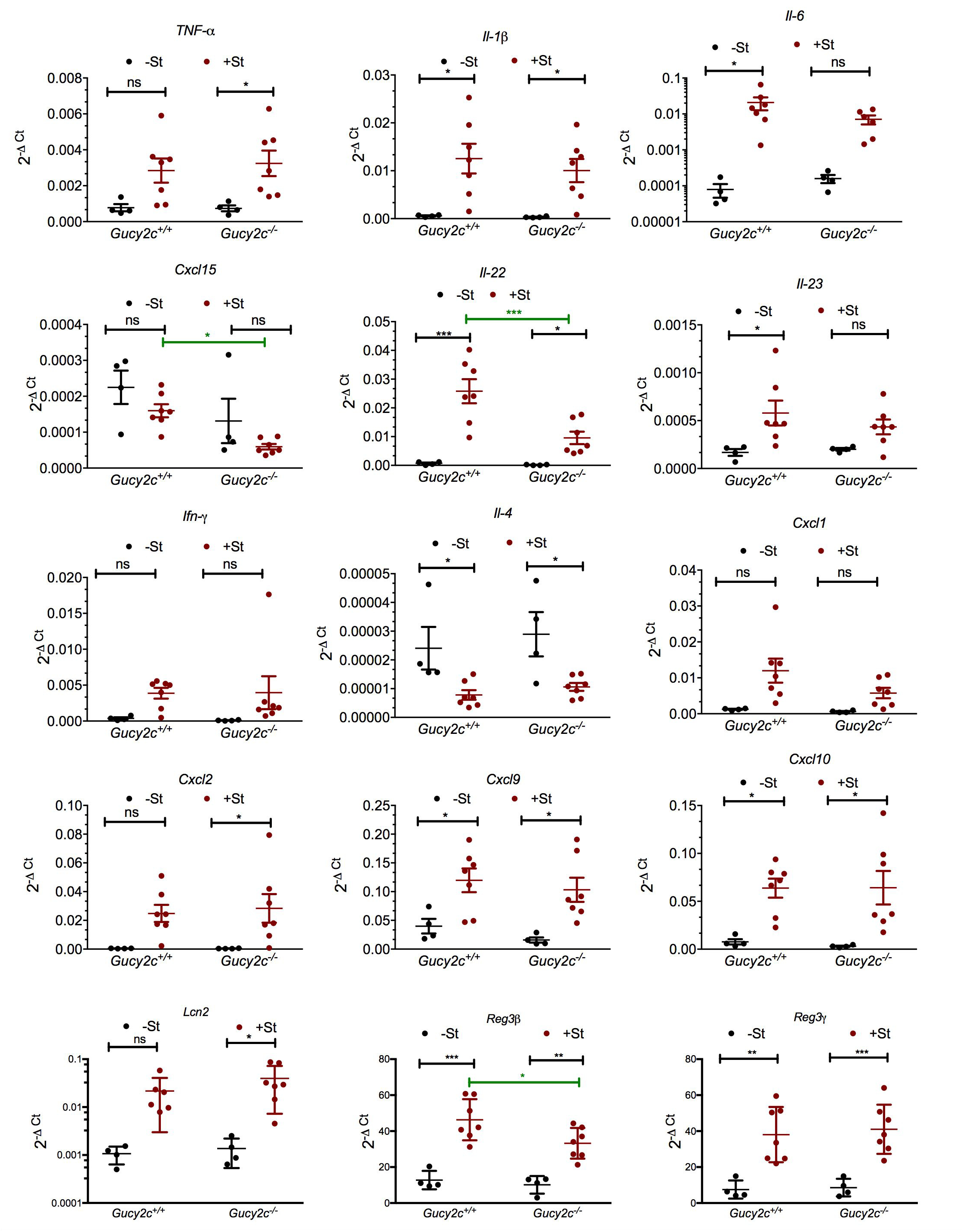
*Gucy2c*^-/-^ mice show reduced transcript levels of IL-23/IL-22 in the ileum following infection. Orally infected *Gucy2c*^+/+^ and *Gucy2c*^-/-^ mice were sacrificed 3 days post infection (+St) along with the control uninfected mice (-St). RNA was isolated from the harvested ileal tissues and quantitative real-time PCR analysis was performed to estimate the transcript levels of *Tnf-α, Il-1b, Il-6, Cxcl-15, Il-22, Il-23, Ifn-γ, Il-4, Cxcl-1, Cxcl-2, Cxcl-9, Cxcl-10, Lcn 2, Reg3β* and *Reg3γ*. The data shown are from individual mice and lines depict the mean ± SD. The experimental groups were analysed for statistical significance using twoway ANOVA, \**p* ≤ 0.05, \*\**p* ≤ 0.01, \*\*\**p* ≤ 0.001 and ns: not significant. Green bars represent values which differed significantly between *Gucy2c*^+/+^ and *Gucy2c*^-/-^ mice.

### Reduced *Lactobacilli* in *Gucy2c*^-/-^ mice may contribute to initial susceptibility of mice to *Salmonella* infection

The gut microbiota is known to influence infection progression by *Salmonella* (17, 18, 25, 26). We therefore monitored the major Phyla of bacteria present in the mouse gut, and also *Lactobacillus sp* which has been reported to protect from *Salmonella* infection (25–28). We prepared genomic DNA from stool samples from both *Gucy2c*^+/+^ and *Gucy2c*^-/-^ mice, and performed real-time PCR using universal and Phyla specific primers for the 16S rRNA genes. As shown in Fig. 6, while the major Phyla (Bacteriodetes and Firmicutes) were largely unchanged in both strains of mice, *Lactobacillus sp* were significantly reduced. Since *Lactobacilli* have been reported to attenuate the severity of salmonellosis in mice (25), we propose that the initial environment of the gut of *Gucy2c*^-/-^ mice could have also contributed to initial infection and uptake of *S.* Typhimurium, culminating in the increased susceptibility of *Gucy2c*^-/-^ mice to infection.

**Figure 6.**
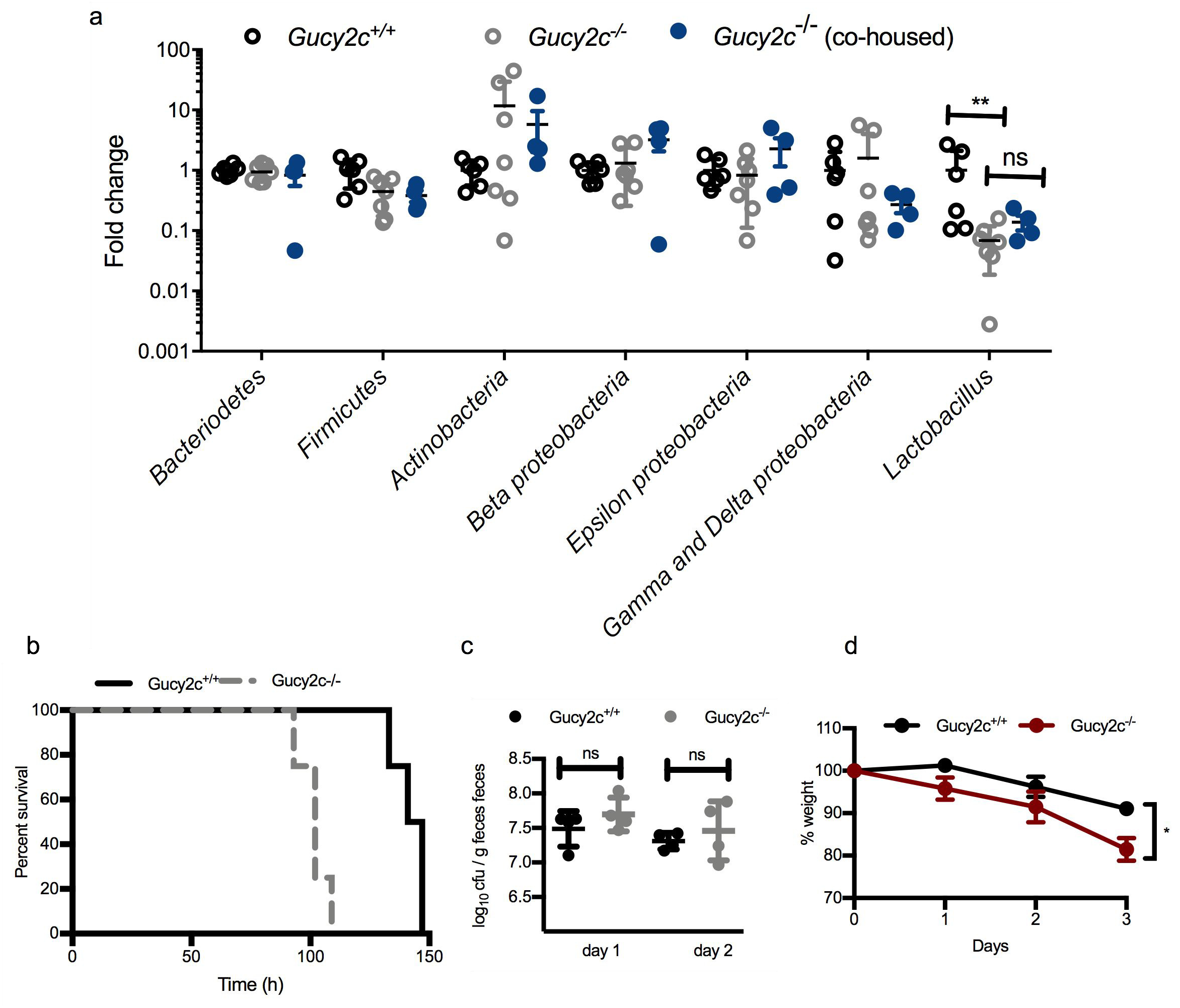
Abundance of major microbial phyla in *Gucy2c*^+/+^ and *Gucy2c*^-/-^ mice. (a) Fecal pellets from uninfected *Gucy2c*^+/+^ and *Gucy2c*^-/-^ mice, and *Gucy2c*^-/-^ mice co-housed with wild type mice, were collected and the bacterial DNA was isolated from the samples. Quantitative real-time PCR was performed to analyse levels of gut microbiota using Phyla specific primers, normalized to universal 16S primers. Values shown represent individual mice and lines show the mean ± SD. Statistical analysis was performed using non-parametric Mann-Whitney *U* test, \*\**p* ≤ 0.01. (b) *Gucy2c*^+/+^ and co-housed *Gucy2c*^-/-^ (4 each) were infected with St and housed together in multiple cages during the course of infection. Survival of mice was checked every 8h. (c) Fecal pellets from individual mice were collected 24 h and 48 h following infection of mice (Gucy2c^+/+^, black circles; *Gucy2c*^-/-^, grey circles) and the bacterial load in the feces on day 1 and 2 following infection were estimated. Data show values for individual mice, with error bars showing the mean ± SD. (c) Individual mice were weighted prior to infection and then every day following infection till day 3 (when *Gucy2c*^-/-^ mice had started to succumb to infection). Data shown is the mean ± SD (n=4).

To ensure that alterations in the microbiome in *Gucy2c*^-/-^ mice were directly a consequence of the absence of GC-C in the gut, we co-housed *Gucy2c*^-/-^ mice at the time of weaning with adult wild type mice for 4 weeks. We then monitored major Phyla in the feces collected from co-housed mice. We continued to observe a similar composition of microbiota after co-housing, including the lower levels of *Lactobacilli* in *Gucy2c*^-/-^ mice (Fig. 6a). We infected these co-housed *Gucy2c*^-/-^ mice along with age-matched wild type mice, housed both strains of mice together in multiple cages following infection, and monitored survival. In agreement with earlier results, we saw dramatically poorer survival of *Gucy2c*^-/-^ mice that were cohoused with infected *Gucy2c*^+/+^ mice following infection (Fig. 6a). Similar fecal bacterial loads (Fig. 6b) were observed in all mice, but *Gucy2c*^-/-^ mice showed a marked reduction in body weight by day 3. Therefore, these co-housing experiments suggest that susceptibility of *Gucy2c*^-/-^ mice to *Salmonella* infection is caused by genetically-determined changes in gut response and microbiome composition.

## Discussion

GC-C-mediated cGMP pathways are essential regulators of intestinal homeostasis, including fluid-ion secretion and cell proliferation(23). The role of GC-C in diarrhea caused by enterotoxigenic *E. coli* that produce heat-stable enterotoxins is well established, since GC-C serves as the receptor for the toxin (11). However, the roles of GC-C, if any, during severe enteric infections, have not been investigated. Here we have utilized the *S.* Typhimurium infection model in mice which results in an acute, progressive and lethal infection, to study the contribution of GC-C during *Salmonella* pathogenesis. Our results show that GC-C provides protection to oral *S.* Typhimurium infection, and absence of GC-C was correlated with reduced ileal damage and changes in the levels of some cytokines and antimicrobial peptides (AMPs) (1, 24) (Fig. 7). Interestingly, the rapid mortality seen in *Gucy2c*^-/-^ mice was not dependent on increased systemic infection, since bacteria colonized extra-intestinal tissue to equivalent extents in both *Gucy2c*^+/+^ and *Gucy2c*^-/-^ mice.

**Figure 7.**
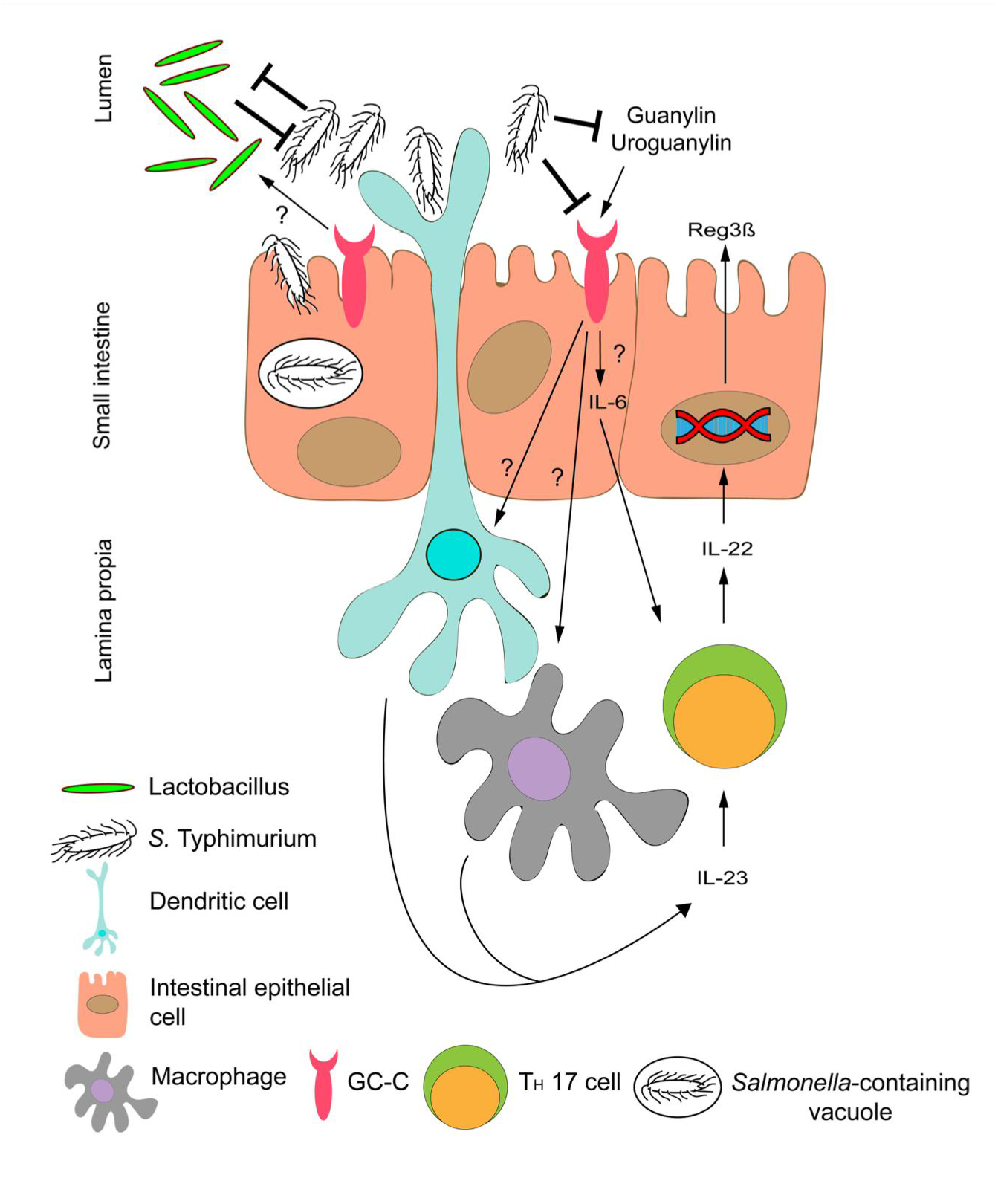
Putative role of GC-C during Salmonella infection. The IL23/IL22 arm of the innate immune response during *Salmonella* infection is regulated by GC-C. Post oral infection in mice, *Salmonella* competes with the commensal *Lactobacillus* to invade the epithelial and dendritic cells of the gut. *Salmonella* downregulates the expression of the GC-C receptor and its endogenous ligands, guanylin and urouanylin. Probable interaction of GC-C with dendritic cells and macrophages along with IL-6 upregulation leads to induction of IL-23 expression, which activates the TH17 cells to express the cytokine IL-22, which in turn induces the expression of the AMP, Reg3β, in epithelial cells.

Glucocorticoids (GCs) are steroid hormones which are secreted by the adrenal glands and possess immunosuppressive properties. GCs are an indicator of stress (29) and are upregulated during *S.* Typhimurium infection in mice (21). At 48 h post infection, serum cortisol levels in *Gucy2c*^-/-^ mice were significantly higher than that seen in *Gucy2c*^+/+^ mice. Thymic atrophy accompanies numerous infections and GCs contribute towards depletion of the developing CD4^+^CD8^+^ immature thymocytes during infections by pathogens such as *Listeria monocytogenes,* type A *Francisella tularensis*, *Trypanosoma cruzi* and *S.* Typhimurium (20, 21). Since *Gucy2c*^-/-^ mice displayed early elevated levels of serum cortisol (Fig. 2a), infection-induced thymic atrophy in *Gucy2c*^-/-^ mice was indeed observed to be significantly higher than in *Gucy2c*^+/+^ mice.

Higher circulating cortisol could be attributed to enhanced colonization and inflammation in *Gucy2c*^-/-^ mice, as evidenced by histopathology of the ileum. *S.* Typhimurium infection results in inflammation of the cecum and colon, with lower inflammation seen in the ileum of mice (30). Germ-free mice display higher ileal load than specific-pathogen-free mice when administered with similar doses of *S.* Typhimurium (31). The higher colonization in the ileum of *Gucy2c*^-/-^ mice may be due to higher translocation or invasion of *Salmonella* in the gut of these mice. Interestingly during Crohn’s disease, selective inflammation in the terminal ileum is observed (32). In addition, the standard therapy for inflammatory bowel disease (IBD) is administration of corticosteroids, which facilitate immunosuppression (33). The elevated levels of cortisol observed in *Gucy2c*^-/-^ mice during infection (Fig. 2a) might be a compensatory mechanism to reduce inflammation of the ileum.

*Salmonella* pathogenesis results in altered gene expression profile in the intestine (1). The transcript levels of GC-C and its endogenous ligands, guanylin and uroguanylin were downregulated upon infection in the ileum (Fig. 4). However, the transcript levels of other downstream effectors of the GC-C pathway such as *PkgII, Cftr and Pde5* were not differentially modulated post infection. NHE3 is an abundantly expressed Na^+^/H^+^ exchanger in the intestine (11, 34). There was a significant increase in the expression of NHE3 upon infection, which occurred independently of the presence of GC-C. The significant increase in NHE3 transcript levels is counterintuitive, since there are reports of downregulation of NHE3 by pro-inflammatory cytokines such as IFN-γ and TNF-α, during EPEC infection and inflammatory bowel disease (34). Mechanisms that lead to transcriptional regulation of NHE3 upon *Salmonella* infection are unknown and warrant further investigation.

There are only two reports of disruption of GC-C signaling at both receptor and ligand levels, as we see here on *Salmonella* infection (Fig. 4). In ulcerative colitis patients, a severity-dependent downregulation in expression of GC-C, guanylin and uroguanylin in the colonic mucosa, is observed (35). GC-C and its ligands are also downregulated in inflamed colonic IBD mucosa of patients, (36) and in 2,4,6-trinitrobenzene sulphonic acid-induced colitis in rats (37). The significant downregulation of GC-C and its ligands upon *Salmonella* infection could therefore occur as a consequence of altered composition of the tissue following the inflammation induced by infection.

*Salmonella* infection and invasion of the intestine induce inflammatory responses including upregulation of pro-inflammatory cytokines (24). The induction of TNF-α in the gastrointestinal tract during *Salmonella* infection is well known (5, 6). Low amounts of TNF-α confer protection to mice towards *Salmonella* infection (38), while administration of higher doses causes extensive histopathological damage, organ dysfunction and death within minutes to hours due to respiratory arrest (39, 40). Following infection, there was no statistical difference between transcript levels of TNF-α in the ileum in wild type and *Gucy2c*^-/-^ mice. TNF-α mediates the depletion of goblet cells during *S.* Typhimurium infection in mice, as pre-treatment with anti-TNFα antibody restores the goblet cell numbers and mucin profiles (6). We however see an increase in goblet cell numbers on infection in the small intestine in *Gucy2c*^-/-^ mice, suggesting a novel means of regulating goblet cell number by GC-C.

A critical arm of the antimicrobial innate immune response is the production of epithelial cell-derived AMPs. This antimicrobial response is mediated via the IL-22 cytokine, which induces the expression of AMPs in the intestinal epithelial cells (1). IL-22 is produced by leukocytes such as T helper type 17 (T_H_17) cells in response to IL-23 produced by infected dendritic cells and macrophages (41) and/or IL-6 produced by intestinal epithelial cells (42). AMPs induced by IL-22 include REG3β, REG3γ, lipocalin 2 and calprotectin. REG3β and REG3γ are bactericidal C-type lectins and eliminate Gram-positive bacteria (1, 43). Lipocalin 2 and calprotectin do not kill bacteria directly but starve them of metals which are essential for their growth (1). This AMP response has been demonstrated to impart a growth advantage to Gram-negative pathogens such as *Salmonella* over commensal bacteria (44). *Reg3γ*^-/-^ and *Reg3β*^-/-^ mice are more susceptible to *Salmonella enteritidis* infection, implying a protective role of these AMPs against this pathogen (45, 46). The lower infection-induced expression of IL-22 in *Gucy2c*^-/-^ mice (Fig. 5) may affect the induction of AMPs in the epithelial cells, and indeed, the transcript levels of Reg3β in *Gucy2c*^-/-^ mice were found to be significantly lower upon infection than in *Gucy2c*^+/+^ mice. This may also partly account for the susceptibility of these mice to *Salmonella* infection.

Innate lymphoid cells are potent producers of IL-22 after intestinal injury. Importantly, IL-22 has been shown to induce intestinal epithelial regeneration, by increasing proliferation of intestinal stem cells (47). *Gucy2c*^-/-^ mice showed a reduction in *Il-22* transcripts on infection (Fig. 6). It is possible that the increased intestinal damage seen in *Gucy2c*^-/-^ mice could be a consequence of reduced proliferation of stem cells and therefore replenishment of epithelial cells.

The gastrointestinal tract is a rich source of aryl hydrocarbon receptor (AhR) ligands (48). Ahr protects the gut upon challenge with pathogenic bacteria (49)by increasing Il-22 expression. Thus, AhR-deficient mice succumb to *Citrobacter rodentium* infection, and ectopic expression of IL-22 protected mice from this early mortality (49). It would therefore be of interest to monitor AhR expression in the gut of *Gucy2c*^-/-^ mice and determine whether this contributes to reduced *Il-22* transcripts in infected mice.

The gut microbiota is a critical determinant for susceptibility to various infectious pathogens (50) including *S.* Typhimurium (17, 18, 25, 26). The gut microbiota can either promote resistance to colonization of pathogens or assist and enhance its virulence (50). In our study, there were no significant differences in the abundance of the major phyla (i.e., Bacteriodetes, Firmicutes, Actinobacteria and Proteobacteria) in *Gucy2c*^+/+^ and *Gucy2c*^-/-^ mice (Fig. 6a). However, we observed a significant decrease in the abundance of *Lactobacillus* in *Gucy2c*^-/-^ mice, as has been reported earlier (16). We observed this decrease even following cohousing of *Gucy2c*^-/-^ mice with wild type mice. (Fig. 6). Different species and isolates of *Lactobacillus* have been shown to exert antagonistic activity on Salmonella invasion and colonization (25–28). For example, *Lactobacillus acidophilus* inhibits adhesion, invasion and dissemination of attenuated *S.* Typhimurium in BALB/c mice (25). In vitro studies in human intestinal Caco-2 cells reveal that *Lactobacillus acidophilus* attenuates *Salmonella*-induced intestinal inflammation via transforming growth factor β/MIR21 signaling pathway (26). The lower abundance of specific *Lactobacillus sp.* might inherently predispose *Gucy2c*^-/-^ mice to *Salmonella* infection, and contribute to the higher degree of translocation and epithelial damage in the ileum.

Microbiota dysbiosis has been reported in mice null for genes involved in fluid-ion homeostasis of the gut (51, 52). It is possible that GC-C enhances growth and colonization of *Lactobacillus sp.* in the gut by providing and maintaining the luminal pH and/or electrolyte distribution and balance. Certain species of *Lactobacillus* have been shown to produce indole-3-aldehyde that is an Ahr ligand. (49). Metagenomic analysis of the microbiome of *Gucy2c*^-/-^ mice may reveal another mechanism by with reduced levels of specific *Lactobacilli* could affect IL-22 production in an AhR-dependent manner.

In summary, we have studied the role of GC-C during *S.* Typhimurium infection in mice and have identified GC-C as a crucial host factor that provides protection against gut-mediated infection by *S.* Typhimurium. Our results, summarized in Fig. 7, indicate that *Salmonella* infection results in downregulation of GC-C ligands and a modest but significant downregulation of GC-C. Absence of GC-C in the gut modulates the innate immune response to *Salmonella* and cytokine production, possibly affecting neutrophil migration and/or stem cell proliferation in the distal ileum. We speculate, therefore, that administration of GC-C ligand analogs may alleviate *Salmonella*-mediated symptoms and pathology. Linaclotide is a FDA approved oral GC-C ligand and is used in treatment of constipation and chronic idiopathic constipation. This drug is effective against visceral pain associated with constipation predominant inflammatory bowel syndrome (C-IBS) (53). Therefore, paradoxically, a target for a bacterial toxin that causes diarrhoea may act as a therapeutic target for treatment and alleviation of *Salmonella*-mediated intestinal symptoms and pathology.

## Materials and Methods

### Bacterial cultures

The *S.* Typhimurium NCTC 12023 strain was used for mice infections. A single isolated colony of *S.* Typhimurium grown on a *Salmonella-Shigella* (SS) agar plate was used to grow culture the pre-inoculum. The overnight pre-inoculum was used at 0.2% in 50 mL of Luria broth and cultured for 3 hours at 37°C and 160 rpm to obtain a log phase culture. The bacterial culture was washed and resuspended in sterile phosphate-buffered saline (PBS) and used for infection (21).

### Mice

*Gucy2c*^-/-^ obtained from the Jackson laboratory, were backcrossed with C57BL/6 mice for more than 10 generations with multiple founder wild type mice. *Gucy2c*^+/+^ and *Gucy2c*^-/-^ mice were bred in the same vivarium, as described previously (15), and housed in a clean air facility in multiple cages, separated based on sex and strain. Temperature (22 ± 2°C) and humidity (55 ± 10%) were maintained with a 12 h light/dark cycle. Mice had access to laboratory chow and water *ad libitum.* Chow was procured from Rayans Biotech, Hyderabad, India, and contained ~ 24% protein, 6% oil, and 3% dietary fiber. Mice aged 6-8 weeks of either sex, weighing 18-30 grams were used for experiments, and following infection housed in multiple cages to rule out cage-dependent effects.

In some experiments, *Gucy2c*^-/-^ mice, at the time of weaning, were co-housed with adult wild type mice for 4 weeks in multiple cages. Feces were collected from co-housed *Gucy2c*^-/-^ mice, and mice then placed in a cage along with the wild type mice to be used for infection. Three days later, mice were used for infection.

### Ethics statement

The experiments were performed in agreement with the Control and Supervision rules, 1998 of Ministry of Environment and Forest Act (Government of India) and the Institutional Animal Ethics Committee of the Indian Institute of Science (IISc). *Gucy2c*^+/+^ and *Gucy2c*^-/-^ mice were bred and maintained at the Central Animal Facility of IISc (Registration number: 48/1999/CPCSEA, dated 1/3/1999). The experimental protocols were approved by the ‘Committee for Purpose and Control and Supervision of Experiments on Animals’ (CPCSEA), permit number: CAF/Ethics/216/2011. The details of the national guidelines can be found on: http://envfor.nic.in/division/committee-purpose-control-and-supervision-experiments-animals-cpcsea (21).

### Mice infection

The mice were infected with *S.* Typhimurium either orally or intra-peritoneally and survival was monitored. For infections via the intra-peritoneal route, ~750 bacteria/mouse were administered, while for oral infection, ~10^8^ bacteria/mouse were used. The bacterial culture was resuspended in PBS and 0.5 mL was administered either intraperitoneally or by oral gavage.

Fresh fecal samples were collected following infection, weighed, homogenized in PBS and the appropriate dilutions were plated on SS agar plates. For quantification of CFU burden in organs, the mice were sacrificed at the indicated time points and the organs were harvested, weighed and homogenized in PBS. Appropriate dilutions were plated on SS agar plates (20).

### Serum cytokines and cortisol estimation

Serum TNF-α, IL-6 and IFN-γ were quantified using ELISA kits (eBioscience, USA), while serum cortisol amounts were measured using the AccuBind ELISA kit (Monobind Inc., USA) according to the manufacturer’s instructions (21).

### Flow cytometric analysis of thymocytes

On indicated days, uninfected and infected mice were sacrificed, thymi dissected and washed in PBS. The organs were disrupted with a pair of forceps and the cell suspensions were passed through a fine wire mesh to obtain a single cell suspension, and viability of the cells was estimated by Trypan blue exclusion assay using a haemocytometer. Thymocytes were stained for cell surface expression of CD4 and CD8 to estimate the major cell subpopulations. Anti-mouse CD4-APC (17-0041-83) and anti-mouse CD8-PE (12-0081-85) were purchased from eBioscience. Cells were incubated with fluorochrome-conjugated antibodies at 4°C for 45 mins. Subsequently, the cells were washed twice with PBS and fixed in 0.5% paraformaldehyde. The cells were acquired on the FACSVerse^™^ flow cytometer (BD Bioscience USA). Baseline instrument application and compensation settings were set using unstained and single stained samples respectively. Single events were gated on the basis of forward scatter-area versus forward scatter-height, while live events were gated on the basis of forward scatter-area versus side scatter-area. These selections were considered to exclusively analyse single live events. CD4 versus CD8 density plots were constructed for thymocytes and the major cell populations were quantified. WinMDI and FlowJo (9.8.5) software were used to plot and analyse the results (21).

### Histological analysis

Histological analysis was performed on ileal sections, which were fixed in 4% paraformaldehyde in PBS (pH 6.9) and Periodic Acid Schiff (PAS) staining was performed (48). Briefly, following fixation, sections were dehydrated by serial immersion in increasing concentrations of ethanol and finally in paraffin. The tissues were embedded in paraffin and 4 μm sections were obtained using a microtome (Leica Biosystems, Germany). The sections were dewaxed and rehydrated by serial immersion in decreasing concentrations of ethanol. Sections were stained with PAS (Avinash Chemicals, Bangalore, India) following the manufacturer’s instructions. The nuclei were subsequently stained with hematoxylin (Sigma-Aldrich, USA). Sections were then dehydrated and mounted. The sections were observed using an Olympus XI81 microscope (Olympus Corporation Tokyo, Japan).

### RNA isolation and quantitative real time PCR analysis

A portion of the distal ileum was isolated and stored in TRI reagent (RNAiso Plus, TaKaRa). The RNA was isolated using the QIAGEN RNeasy kit, according to manufacturer’s protocol and 4 μg of RNA was reverse transcribed to cDNA using Revert Aid reverse transcriptase (Thermo-Scientific, USA). Real-time PCR was performed using SYBR^®^ Premix Ex Taq^™^ (Tli RNaseH Plus) on a CFX96 Touch^™^ Real-Time PCR Detection System (BIO-RAD, USA). Three housekeeping genes Glyceraldehyde 3-phosphate dehydrogenase (*Gapdh*), TATA box binding protein (*Tbp*) and β-actin (*Actb*) were used for internal normalization controls. Since all three genes showed equivalent results, *Gapdh* was used for subsequent normalization of the real time PCR data. Data is expressed as 2^-ΔCT^. The sequences of primers used for quantitative real-time PCR were obtained from those validated at the PrimerBank database and are available on request (54–56).

### Quantitative real-time PCR amplification of 16S sequences of fecal microbiota

Bacterial DNA was isolated from fecal pellets using the QIAamp^®^ DNA Stool Mini Kit according to manufacturer’s instructions. The abundance of total bacteria and specific intestinal bacterial phylum, class and species was quantified by real-time PCR analysis by SYBR^®^ Premix Ex Taq^™^ (Tli RNaseH Plus) on a StepOnePlus real-time PCR machine (Applied Biosystems, USA) using 16S primers published earlier (57, 58). The efficiencies of each primer were estimated (data not shown) and were found to be > 90%. Relative bacterial abundance was determined using the ΔΔCt method by employing universal primers for 16s rDNA for normalization (59). The mean of ΔΔCt values obtained for each Phyla seen in wild type mice was calculated, and this value was used to express the fold-change in individual ΔΔCt values for wild type and *Gucy2c*^-/-^ mice, to obtain the data shown in Fig. 6.

### Statistical analysis

All data was analysed using GraphPad Prism 7. The pooled results from independent experiments are depicted as mean ± SD. All data was tested for normality distribution by the Shapiro-Wilk normality test and found to be normally distributed. Statistical significance among groups of mice were determined using two-way ANOVA, with multiple comparisons, and FDR by the two-stage linear step-up procedure of Benjamini, Krieger and Yekutieli. Students t-test with 95% confidence interval was used for comparing abundance of microbiota in fecal samples. Statistical significance between the mice survival curves were obtained by the Gehan-Breslow-Wilcoxon test. The denoted p values are as follows: \**p*<0.05, \*\**p*<0.01, \*\*\**p*<0.001.

## Acknowledgements

The authors acknowledge facilities provided by the Central Animal Facility at the Indian Institute of Science for housing mice. Financial support from the Department of Biotechnology (DBT), Government of India (SSV) is acknowledged, as well as support provided to the Division of Biological Sciences in the form of Grant-In-Aid by DBT. SSV is a JC Bose National Fellow and support from the Department of Science and Technology, Government of India, is acknowledged. SSV is a recipient of a Royal Society International Collaboration Award for Research Professors along with Prof. Gad Frankel from Imperial College London. SSV would like to thank Prof. Gad Frankel and Dr. Avinash R. Shenoy in the Centre for Molecular Bacteriology and Infection at Imperial College, London, for useful discussions.

